# Adjusting for principal components can induce spurious associations in genome-wide association studies in admixed populations

**DOI:** 10.1101/2024.04.02.587682

**Authors:** Kelsey E. Grinde, Brian L. Browning, Alexander P. Reiner, Timothy A. Thornton, Sharon R. Browning

## Abstract

Principal component analysis (PCA) is widely used to control for population structure in genome-wide association studies (GWAS). Top principal components (PCs) typically reflect population structure, but challenges arise in deciding how many PCs are needed and ensuring that PCs do not capture other artifacts such as regions with atypical linkage disequilibrium (LD). In response to the latter, many groups suggest performing LD pruning or excluding known high LD regions prior to PCA. However, these suggestions are not universally implemented and the implications for GWAS are not fully understood, especially in the context of admixed populations. In this paper, we investigate the impact of pre-processing and the number of PCs included in GWAS models in African American samples from the Women’s Women’s Health Initiative SNP Health Association Resource and two Trans-Omics for Precision Medicine Whole Genome Sequencing Project contributing studies (Jackson Heart Study and Genetic Epidemiology of Chronic Obstructive Pulmonary Disease Study). In all three samples, we find the first PC is highly correlated with genome-wide ancestry whereas later PCs often capture local genomic features. The pattern of which, and how many, genetic variants are highly correlated with individual PCs differs from what has been observed in prior studies focused on European populations and leads to distinct downstream consequences: adjusting for such PCs yields biased effect size estimates and elevated rates of spurious associations due to the phenomenon of collider bias. Excluding high LD regions identified in previous studies does not resolve these issues. LD pruning proves more effective, but the optimal choice of thresholds varies across datasets. Altogether, our work highlights unique issues that arise when using PCA to control for ancestral heterogeneity in admixed populations and demonstrates the importance of careful pre-processing and diagnostics to ensure that PCs capturing multiple local genomic features are not included in GWAS models.

**Author Summary:** Principal component analysis (PCA) is a widely used technique in human genetics research. One of its most frequent applications is in the context of genetic association studies, wherein researchers use PCA to infer, and then adjust for, the genetic ancestry of study participants. Although a powerful approach, prior work has shown that PCA sometimes captures other features or data quality issues, and pre-processing steps have been suggested to address these concerns. However, the utility and downstream implications of this recommended preprocessing are not fully understood, nor are these steps universally implemented. Moreover, the vast majority of prior work in this area was conducted in studies that exclusively included individuals of European ancestry. Here, we revisit this work in the context of admixed populations—populations with diverse, mixed ancestry that have been largely underrepresented in genetics research to date. We demonstrate the unique concerns that can arise in this context and illustrate the detrimental effects that including principal components in genetic association study models can have when not implemented carefully. Altogether, we hope our work serves as a reminder of the care that must be taken—including careful pre-processing, diagnostics, and modeling choices—when implementing PCA in admixed populations and beyond.

## 1 Introduction

There is considerable variability in global ancestry—the genome-wide proportion of genetic material inherited from each ancestral population—across individuals in admixed populations [1–5]. Heterogeneous global ancestry, as with other types of population structure, can lead to spurious associations in genome-wide association studies (GWAS) [6–9]. In fact, some authors have cited the ancestral heterogeneity of admixed populations, and the statistical challenges it poses, as one reason why these populations have been historically underrepresented in GWAS [10–14]. Spurious associations can arise in GWAS in ancestrally heterogeneous populations when global ancestry confounds the association between genotypes and the phenotype of interest. This confounding occurs when the genetic variant being tested differs in frequency across ancestral populations (i.e., global ancestry is associated with genotype) and global ancestry also is correlated with the phenotype of interest via, for example, environmental factors or causal loci elsewhere in the genome that differ in frequency across ancestral groups.

A number of methods for detecting and controlling for ancestral heterogeneity in genetic association studies have been proposed. Early approaches included restricting analyses to subsets of ancestrally homogeneous individuals [15], performing a genome-wide correction for test statistic inflation due to ancestral heterogeneity via genomic control [6], and using family-based designs [16]. More recently, approaches based on mixed models have been proposed [17–19], using random effects to control for both close (e.g., due to family-based sampling) and distant (e.g., due to shared ancestry) relatedness across individuals. When studies do not include closely related individuals, a simpler approach is to include inferred global ancestry as a fixed effect in marginal regression models [7, 20]. This fixed effects adjustment for global ancestry is used extensively in published studies (e.g., [21–28]), with global ancestry inferred using either model-based ancestry inference methods or principal component analysis.

Model-based approaches for ancestry inference (e.g., frappe [29], STRUCTURE [30], ADMIXTUR [31], RFMix [32], FLARE [33]) model the probability of observed genotypes given unobserved ancestry and allele frequencies in each ancestral population. These approaches can be used to estimate global ancestry proportions, also known as admixture proportions, which can then be included as covariates in GWAS models to adjust for ancestral heterogeneity. One of the challenges of using these model-based approaches to infer global ancestry is that the total number of ancestral populations usually needs to be pre-specified. In addition, many of these approaches are supervised, requiring reference panel data from each ancestral population of interest to estimate allele frequencies. Furthermore, ancestry inference is typically conducted at a continental level (e.g., African versus European). Finer levels of population structure could thus be missed, although some recent efforts have considered global ancestry inference on a sub-continental scale [34–38]. Recent discussion has also called attention to the potential harms that may be caused, even if inadvertently, by using continental ancestry as a population descriptor in human genetics research [39].

In many ways, principal component analysis (PCA) provides an attractive alternative to model-based genetic ancestry inference. PCA is an unsupervised approach for inferring global ancestry that does not require reference panel data or pre-specification of the number of ancestral populations of interest, and it is capable of capturing sub-continental structure [23]. To infer global ancestry using PCA, we preform a singular value decomposition of the matrix of standardized genotypes (i.e., **X** = **UDV**^T^) or, equivalently, an eigenvalue decomposition of the genetic relationship matrix (i.e., **XX***^T^* = **UD**^2^**U**^T^). The top principal components (PCs), **u**_1_, **u**_2_*, . . .*, tend to reflect global ancestry [40, 41]. This process is implemented by a variety of software packages (e.g., EIGENSTRAT [7], SNPRelate [42], PC-AiR [43]).

To adjust for ancestral heterogeneity in GWAS, researchers must choose some number of PCs to include as covariates in GWAS regression models. Determining this number of PCs needed to capture global ancestry is non-trivial. Numerous techniques have been proposed, including formal significance tests based on Tracy-Widom theory [7, 40], examining inflation factors [5, 44] or the proportion of variance explained by each PC [5, 44, 45], comparing PCs to self-reported race/ethnicity [5], and keeping PCs that are significantly associated with the trait [24, 46]. Often, the number of PCs selected is on the order of one to ten [47], but in practice it is also common to see applications in which many more PCs are used—in some cases, more than may be necessary to capture global ancestry. The latter has been justified by prior work suggesting that including higher-order PCs can provide the safeguard of removing “virtually all stratification” [48] at the cost of perhaps only “subtle” decreases in power [49].

Another challenge in using PCA to control for ancestral heterogeneity lies in ensuring that PCs actually reflect global ancestry. PCs may instead capture relatedness across samples [9, 40, 43, 50], array artifacts or other data quality issues [7, 9, 40, 51]. Numerous studies have also shown that PCs can capture small regions of the genome with extensive, long-ranging, or otherwise unusual patterns of linkage disequilibrium (LD) [7, 9, 21, 40, 50–56]. These unusual patterns of LD may result from positive selection [57–59] or other reasons, such as the presence of an inversion [23, 52–54]. When PCs capture these local genomic features, they typically are not as highly correlated with global ancestry or geography, and thus adjusting for such PCs in GWAS may not be as effective at reducing spurious associations caused by population structure [45, 47, 50, 54, 55, 59–61]. Prior studies have also suggested that adjusting for PCs capturing local genomic features may reduce power, particularly if a causal variant is located in one of the regions captured by a PC included in the GWAS model [45, 54–56, 62, 63], or lead to biased GWAS summary statistics [38]. To avoid these issues, many authors have suggested that LD must be addressed when conducting PCA [21–23, 40, 51, 54, 56, 59]. The most commonly implemented technique involves running PCA on a reduced subset of variants after first performing LD pruning, using a program such as PLINK [64] to remove variants that are in “high” LD (e.g., pairwise-correlation *r*^2^ *>* 0.2) with nearby variants [5, 21–23, 43, 44, 46, 50, 51, 55, 59, 60, 63, 65, 66], and/or excluding regions of the genome that are known to have unusual patterns of LD [5, 21–23, 45, 51, 53, 66]. We provide a list of these previously-identified high LD regions and references advocating for their exclusion in Table 1.

**Table 1:** Regions of the genome with high, long-range, or otherwise unusual LD that are often recommended for exclusion prior to running PCA. Start and end physical (base pair) positions are provided with respect to genome build 38. This list is also available for download in builds 36, 37, and 38 at https://github.com/kegrinde/PCA/.

| Chr | Start (bp) | End (bp) | References |
| --- | --- | --- | --- |
| 1 | 47761741 | 51822307 | [51, 53, 66] |
| 2 | 129125957 | 139525961 | [5, 23, 45, 51, 53, 61] |
| 2 | 182309767 | 189427029 | [51, 53, 66] |
| 3 | 47483506 | 49987563 | [51, 53, 66] |
| 3 | 83368159 | 86868160 | [51, 53, 66] |
| 3 | 161899518 | 163699518 | [61] |
| 5 | 98636396 | 101136397 | [51, 53] |
| 5 | 129636408 | 132636409 | [51, 53, 66] |
| 5 | 136136412 | 139136412 | [51, 53] |
| 6 | 23691793 | 38924246 | [5, 22, 23, 45, 51, 53, 61, 66] |
| 6 | 139637170 | 142137170 | [51, 53, 66] |
| 8 | 6455071 | 13598120 | [5, 22, 23, 45, 51–53, 61, 66] |
| 8 | 110918595 | 113918595 | [51, 53, 66] |
| 11 | 88127184 | 91127184 | [51, 53, 66] |
| 12 | 110577812 | 113099475 | [51, 53] |
| 14 | 47061047 | 47961047 | [61] |
| 17 | 42394456 | 46567318 | [5, 23] |
| 20 | 33948533 | 36438183 | [51, 53, 66] |

Despite having been recommended by many authors, the steps of LD pruning and removing known high LD regions (Table 1) prior to PCA are not universally practiced. Reviewing four recent issues of a top human genetics journal, for example, we identified twelve articles that used PCA to control for population structure in genetic association studies; only one of these papers made any mention of having implemented these pre-processing steps. Perhaps one reason for the lack of consistent implementation, or documentation, of these pre-processing steps is a lack of clarity regarding their effectiveness in alleviating the issues posed by PCs capturing local genomic features. For example, past studies provide conflicting evidence regarding the effect of pre-processing on how well PCs capture population structure: LD pruning, or accounting for LD in some other way, has resulted in PCs that better capture population structure in many datasets [50, 54, 59], but not all [7, 50]. Furthermore, while some studies show that models adjusting for PCs generated after LD pruning yield a lower inflation factor [50, 60] or family-wise error rate [54] compared to those that did not, other studies suggest that inflation factors may not be strongly impacted by the set of PCs used [62] or may even be *higher* after LD pruning [45]. As recently noted by Prive et al. [56],“more work is needed to understand these fundamental problems, and to provide precise guidelines for conducting successful GWAS, heritability and other genomic analyses where PCA is used.”

Although it is widely understood that PCs may capture local genomic features, the downstream implications of adjusting for such PCs are not fully understood, the recommended solutions of LD pruning and removing known high LD regions are not universally practiced, and the evidence regarding the effectiveness of these pre-processing steps is not clear cut. Furthermore, the vast majority of the prior work on this topic was conducted in populations of European ancestry [38, 45, 50, 54, 59–62]. Those studies that did consider non-European populations focused on large worldwide, trans-ancestry samples [50, 56]. Recommendations on how best to implement PC-based adjustment for ancestral heterogeneity in admixed populations are lacking. Given that LD patterns differ in admixed populations compared to these other groups, it is plausible that the extent to which PCs capture local genomic features, the downstream implications for GWAS, and even the ideal solutions for these populations may also differ.

This paper seeks to address the open questions pertaining to using PCA to control for ancestral heterogeneity in GWAS in admixed populations. Specifically, we investigate the impact of LD filtering and pruning, as well as choices of the number of PCs to include in analyses, when performing GWAS in admixed populations. Our analyses focus on three samples of unrelated African American individuals from the Women’s Health Initiative SNP Health Association Resource (WHI SHARe) and two contributing studies to the Trans-Omics for Precision Medicine (TOPMed) Whole Genome Sequencing Project: the Jackson Heart Study (JHS) and the Genetic Epidemiology of Chronic Obstructive Pulmonary Disease Study (COPDGene). For comparison, we also consider a subset of unrelated European American individuals from the COPDGene study. First, we investigate the extent to which top PCs capture genome-wide ancestry versus local genomic features, depending on the pre-processing choices made. Then, we conduct simulation studies and provide theoretical results to demonstrate the downstream implications of including PCs that capture local genomic features in GWAS models. Throughout, we explore the extent to which LD filtering and pruning can alleviate these issues. To conclude, we provide suggestions regarding best practices for controlling for ancestral heterogeneity when conducting GWAS in admixed populations.

## 2 Results

### 2.1 Ancestral heterogeneity in WHI SHARe, JHS, and COPDGene African Americans

We focus our analyses on three examples of ancestrally heterogeneous admixed populations. Specifically, we consider samples of unrelated African American individuals from the WHI SHARe, JHS, and COPDGene studies. (Note that these studies also include related individuals, Hispanic Americans, and/or European Americans. To facilitate comparisons, we primarily focus our attention on the subset of mutually unrelated African Americans within each study. Methods to identify these subsets can be found in Section 4.1.) Before investigating the performance of PCA in these samples, we first explored the extent of ancestral heterogeneity present in each. In each sample, we estimated genome-wide admixture proportions using model-based genetic ancestry inference approaches. Inferred admixture proportions for these three samples of African American individuals are presented in Figure 1.

**Figure 1:**
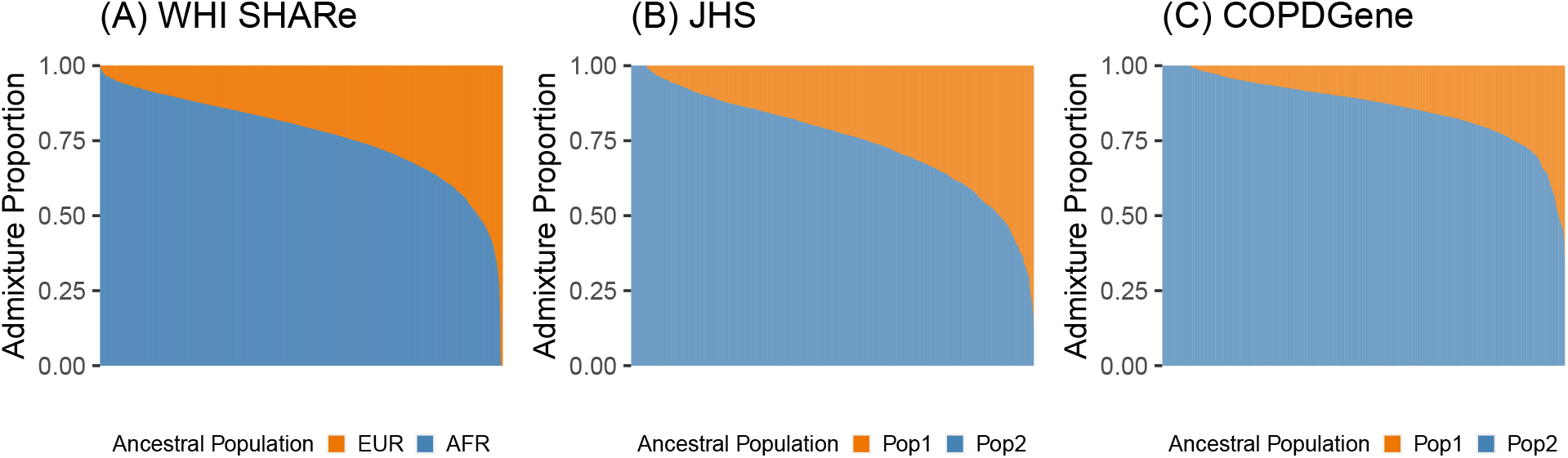
Estimated admixture proportions in three samples of unrelated African American individuals. Each individual is represented by a narrow vertical bar in the plot, and the individuals are sorted by their estimated proportion of African ancestry. The admixture proportions shown here were estimated using RFMix in WHI SHARe (A) and an unsupervised ADMIXTURE analysis in TOPMed JHS (B) and COPDGene (C).

In WHI SHARe, we compared admixture proportion estimates from three techniques. Figure 1A presents admixture proportions estimated as genome-wide average local ancestry, using local ancestry calls from RFMix with two ancestral populations. The reference panel for local ancestry inference consisted of Utah residents with Northern and Western European ancestry (denoted as EUR in Figure 1) and Yoruba in Ibadan, Nigeria (AFR in Figure 1) from the International HapMap Project [67]). These local ancestry based admixture proportion estimates were highly correlated (Pearson correlation *>* 0.998) with estimated admixture proportions from both supervised and unsupervised ADMIXTURE analyses with two ancestral populations.

In TOPMed samples, we performed unsupervised ADMIXTURE analyses with two ancestral populations. Figures 1B and 1C present results for unrelated African American individuals in JHS and COPDGene, respectively. As these analyses were unsupervised, we cannot be certain which ancestral populations are represented and thus we denote the two populations simply as “Pop 1” and “Pop 2” in Figure 1. However, in comparison to the distribution of admixture proportions seen in WHI SHARe African Americans (Figure 1A) and results from prior studies of admixture in African Americans, we hypothesize that “Pop 1” may correspond to European ancestry and “Pop 2” to African ancestry.

In all three samples, we observed considerable variability in the relative proportions of African and European ancestry across individuals. Thus, these three samples exemplify contexts in which careful adjustment for global ancestry will be necessary when performing GWAS.

### 2.2 Top PC correlates strongly with global ancestry

In addition to model-based global ancestry inference, we also performed PCA in the three admixed samples and compared results from the two techniques. We considered sets of PCs computed on the full set of variants that passed initial quality control filters, as well as smaller subsets of variants that remained after LD-based exclusions and/or pruning. Methodological details are provided in Section 4.2.2.

In an African American population, we might expect that only one PC is needed to capture ancestral heterogeneity, at least with respect to differences in the relative proportion of African and European continental ancestry. Comparing estimated admixture proportions to PCs in WHI SHARe, JHS, and COPDGene confirmed this hypothesis. In all three samples of African Americans, the first PC was highly correlated with the inferred proportion of African ancestry (Pearson correlation: 0.993–0.998). Later PCs, however, showed very little correlation with genome-wide continental ancestry (Pearson correlation: -0.015–0.034). We observed similar patterns of correlation between PCs and inferred admixture proportions regardless of the type of LD filtering, or lack thereof, performed prior to running PCA (Supplemental Information Section S3).

### 2.3 Multiple local genomic features captured by later PCs

In these three African American samples, the first PC was highly correlated with global ancestry, whereas later PCs were not. It is possible that these higher-order PCs may reflect sub-continental structure that is not captured by the estimated admixture proportions. However, in many cases we saw that these later PCs captured local genomic features rather than genome-wide ancestry. This was evident from inspection of SNP loadings—which rep-resent the contribution of each variant to each PC—or, similarly, from investigation of the correlation between PC scores and genotypes.

Figure 2 presents the correlation between PCs and genotypes in JHS and COPDGene African Americans when PCs were generated without any prior LD-based pruning or filtering. We see that variants across the genome contributed relatively equally to the first PC, whereas the second, third, and fourth PCs had out-sized contributions from variants on a select number of chromosomes. In JHS, for example, the second PC was particularly highly correlated with variants on chromosomes 6 and 8, and to a lesser extent with variants on chromosomes 2, 3, and 11. We saw similar patterns, although with peaks on different combinations of chromosomes, in COPDGene (Figure 2B) and WHI SHARe African Americans (Figure 3A). The peaks in these genotype-PC correlation plots indicate that those PCs primarily captured variation at these positions rather than genome-wide global ancestry.

**Figure 2:**
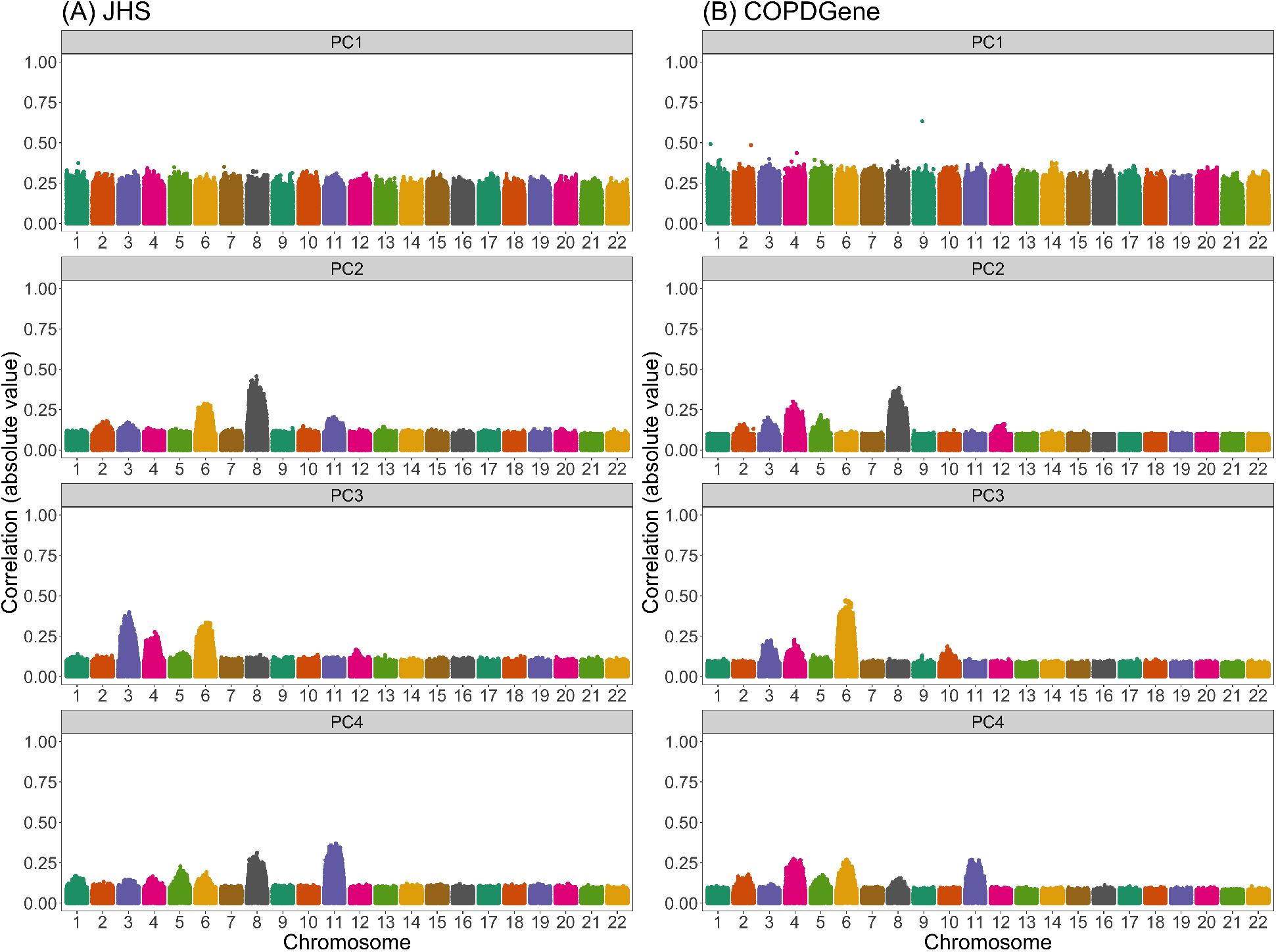
Correlation between naively generated PCs (i.e., PCs that were constructed with-out any prior LD-based filtering or exclusions) and genotypes in JHS and COPDGene African Americans. Each panel plots the absolute value of the correlation between PCs and geno-types (on the y-axis) versus the position along the genome (x-axis). Panels are organized vertically according to which PC is being investigated (1, 2, 3, 4) and horizontally according to the sample (A: JHS, B: COPDGene). Peaks in this plot indicate that a variant has a larger *loading*, i.e., a larger contribution to that PC.

**Figure 3:**
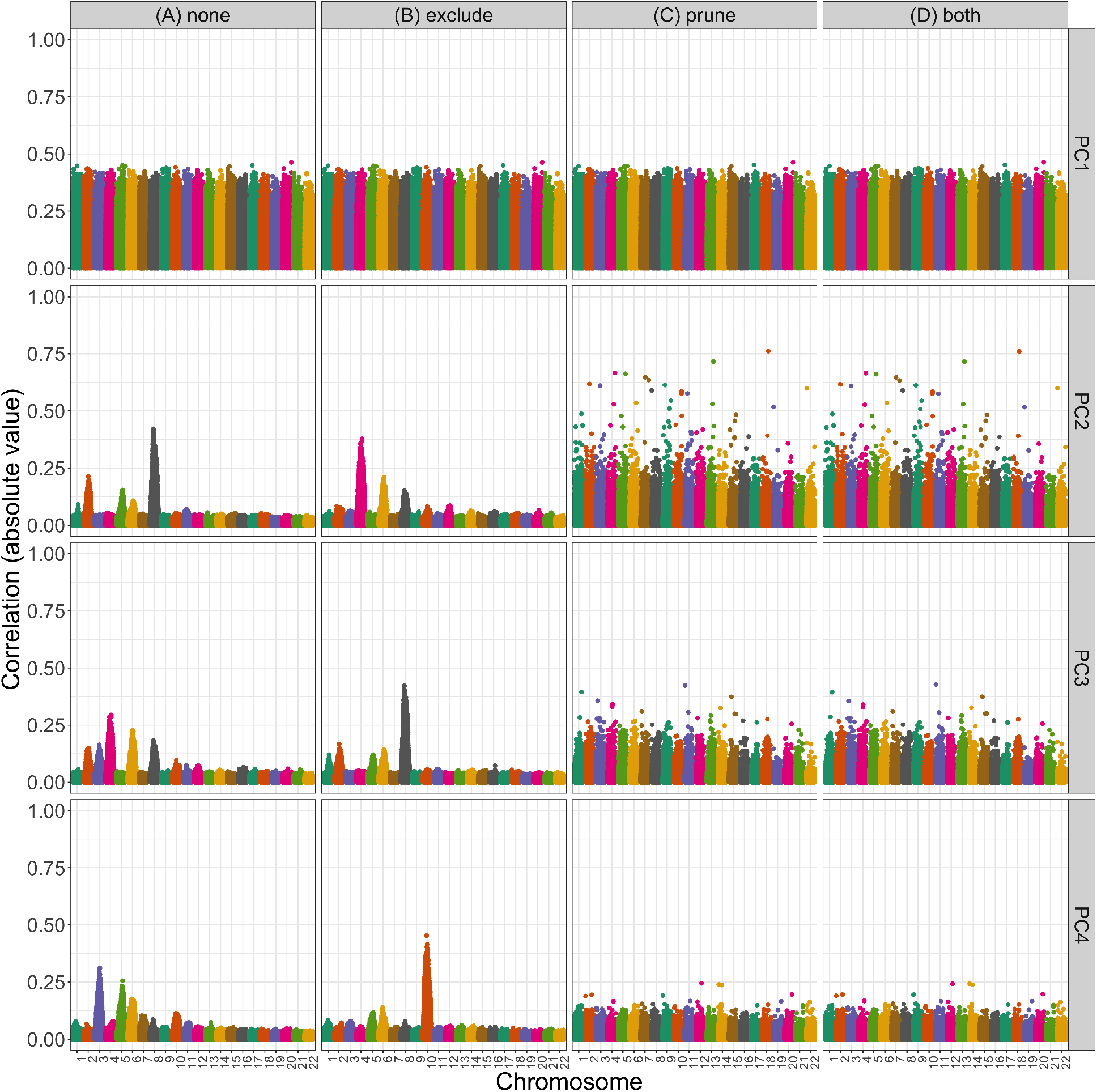
Correlation between PCs and genotypes in WHI SHARe African Americans with different choices of pre-processing. Each panel plots the absolute value of the correlation between PCs and genotypes (on the y-axis) versus the position along the genome (x-axis). Panels are organized vertically according to which PC is being investigated (1, 2, 3, 4) and horizontally according to the level of filtering that was applied prior to running PCA (*none*: all SNPs, *exclude*: after excluding regions in Table 1, *prune*: after LD pruning with an *r*^2^ threshold of 0.1 and window size of 0.5 Mb, and *both*: after both exclusions and LD pruning).

To compare the patterns we saw in these admixed samples to a European population, we also ran PCA in a subset of European American individuals from the COPDGene study. SNP loadings for the first four PCs are presented in Supplemental Figure S9. We see pronounced peaks in the loadings plots particularly for the second and third PC. In both cases, the PCs were highly correlated with variants on a single chromosome (chromosome 11). This pattern of observing a single, pronounced peak in the SNP loadings for a given PC is in agreement with that mentioned in previous studies [38, 45, 50, 54–56, 59–62], but stands in contrast to the pattern of multiple peaks per chromosome that we observed in the JHS, COPDGene, and WHI SHARe African American samples (Figures 2 and 3A). As we will discuss below, this difference in terms of how many regions of the genome are captured by each PC has distinct consequences for GWAS.

### 2.4 Impact of LD pruning on PCA

Previous authors have suggested that this phenomenon of PCs capturing local genomic features arises due to high or otherwise unusual patterns of LD among variants; as a result, they recommend that variants in high LD with one another be removed prior to running PCA. Following these recommendations, we compared the set of PCs based on all variants to PCs generated after first removing regions of the genome known to have high LD (Table 1), performing LD pruning, or both.

Table 2 presents the number of variants that remained after each set of pre-processing steps. Removing previously-identified high LD regions cut the number of variants used for PCA by only about 3% across the three samples we considered. LD pruning, with or without additional Table 1 exclusions, reduced the number of variants more dramatically, removing over 90% of the original variants. Despite the smaller number of variants used for PCA, initial PCs were still very highly correlated with model-based estimates of global ancestry (Figure S13). The greatly reduced set of variants also offered computational advantages for running PCA.

**Table 2:**
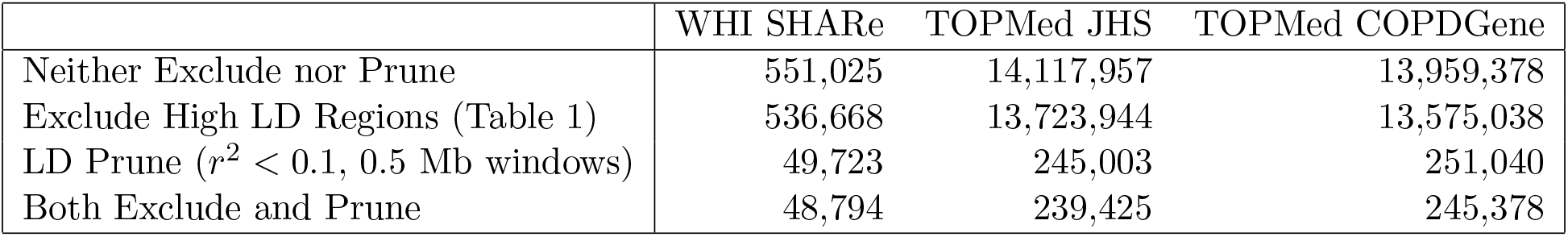
Number of autosomal SNPs remaining after different combinations of pre-processing steps were applied prior to running PCA. Note that TOPMed analyses include only those variants with minor allele frequency greater than 1%.

|  | WHI SHARe | TOPMed JHS | TOPMed COPDGene |
| --- | --- | --- | --- |
| Neither Exclude nor Prune | 551,025 | 14,117,957 | 13,959,378 |
| Exclude High LD Regions (Table 1) | 536,668 | 13,723,944 | 13,575,038 |
| LD Prune ($r^2 < 0.1$, 0.5 Mb windows) | 49,723 | 245,003 | 251,040 |
| Both Exclude and Prune | 48,794 | 239,425 | 245,378 |

Figure 3 illustrates the impact of these pre-processing steps on the correlation between genotypes and PCs in WHI SHARe African Americans. Panel A presents results for PCs that were generated without any prior LD-based filtering or pruning (i.e., using the entire set of 551,025 variants). In this setting, we see that PCs 2–4 captured local genomic features rather than genome-wide ancestry. Panel B shows what happened when we excluded the ≈ 15,000 variants located in the previously-identified high LD regions reported in Table 1: although the pattern of *which* SNPs were driving PCs 2–4 changes, the overarching issue of PCs capturing local genomic features was not resolved. However, after LD pruning with an *r*^2^ threshold of 0.1 and a window size of 0.5 Mb we saw similar patterns with PCs 2–4 as with the first PC: in Panel C, the peaks are gone, with all variants contributing relatively equally to each PC. When we removed previously-identified high LD regions in addition to performing LD pruning (Panel D), we saw very similar patterns of correlation between PCs and genotypes as we did with LD pruning alone (Panel C).

Although LD pruning proved effective in preventing top PCs from capturing local genomic features in the WHI SHARe dataset, careful consideration was need to identify the optimal the choice of LD pruning parameters. When performing LD pruning using a program such as PLINK [64] or SNPRelate [42], two parameters must be selected: *r*^2^ threshold and window size. The *r*^2^ threshold we used in WHI SHARe (*r*^2^ *<* 0.1) is stricter than the default for many software programs and the threshold used in many studies of European populations (*r*^2^ *<* 0.2). When we used the larger *r*^2^ threshold of 0.2 in WHI African Americans, we saw improvement for the second and third PCs, but the fourth continued to capture local genomic features (Supplemental Figure S1). In contrast, LD pruning with the default settings (*r*^2^ *<* 0.2, 0.5 Mb windows) did alleviate issues in the TOPMed COPDGene European American sample (Supplemental Figure S11). In TOPMed JHS African Americans, both the stricter *r*^2^ threshold of 0.1 and a wider window size of 10 Mb were needed (Supplemental Figure S5). Yet, even these strict pre-processing settings were not sufficient in TOPMed COPDGene African Americans: peaks still remained in the PC-genotype correlation plots for the second, third, and fourth PCs after removing regions listed in Table 1 and LD pruning with an *r*^2^ threshold of 0.1 and window size of 10 Mb (Supplemental Figure S7). An even smaller *r*^2^ threshold did not alleviate these issues (Supplemental Figure S8).

### 2.5 Adjusting for PCs that capture local genomic features can induce spurious associations

As we have shown above, top PCs in admixed populations may capture multiple local genomic features, and this can occur even when PCs were generated after first removing previously identified high LD regions (Table 1) and/or performing LD pruning, depending on the parameter settings used. If pre-processing steps are not performed, or if default settings are used and PCs are not carefully checked, GWAS models will then be adjusting for PCs that capture these local genomic features instead of genome-wide ancestry. It remains to be fully understood what the downstream implications would be of adjusting for these PCs in GWAS. We derived expected effect size estimates and conducted simulation studies to investigate these implications further.

Figure 4 presents Manhattan plots from one replicate of our simulation study using real genotype data from WHI SHARe African Americans and simulated traits (see Section 4.3.3 for simulation set-up details). In this particular setting, there was a single causal variant on chromosome 4. We compared results from GWAS models using different ancestral heterogeneity adjustment approaches. As expected, we saw extreme inflation, i.e., statistically significant associations on *every* chromosome, when we did not make any adjustment for ancestral heterogeneity (Figure 4A). When we inferred and adjusted for ancestral heterogeneity using either PCA or estimated admixture proportions, we saw a single peak in our Manhattan plot on chromosome 4—as hoped, given that is where the causal variant was located—with one notable exception. When we adjusted for the first four PCs (as had been done in previous GWAS in WHI SHARe [24, 25]), where those PCs were generated without any prior LD-based pruning or filtering, then we saw a spurious association on chromosome 6 (Figure 4C). However, this spurious association disappeared if we only adjusted for the first of these PCs (Figure 4B). Likewise, no spurious association arose when we adjusted for estimated admixture proportions (Figure 4D) or when we used PCs that were generated after strict LD pruning and Table 1 exclusions (Figure 4, panels E and F). Note that the causal variant, on chromosome 4, and the spurious signal, on chromosome 6, are both located in regions of the genome that are highly correlated with the PCs that were generated without any prior LD pruning (Figure 3). We observed a similar phenomenon in our TOPMed simulations: when GWAS models adjusted for an extraneous, artificially-generated PC that was constructed in such a way that it was highly correlated with the causal variant as well as a variant on a distinct chromosome, a spurious association appeared at that second variant (Supplemental Information Section S5.2).

**Figure 4:**
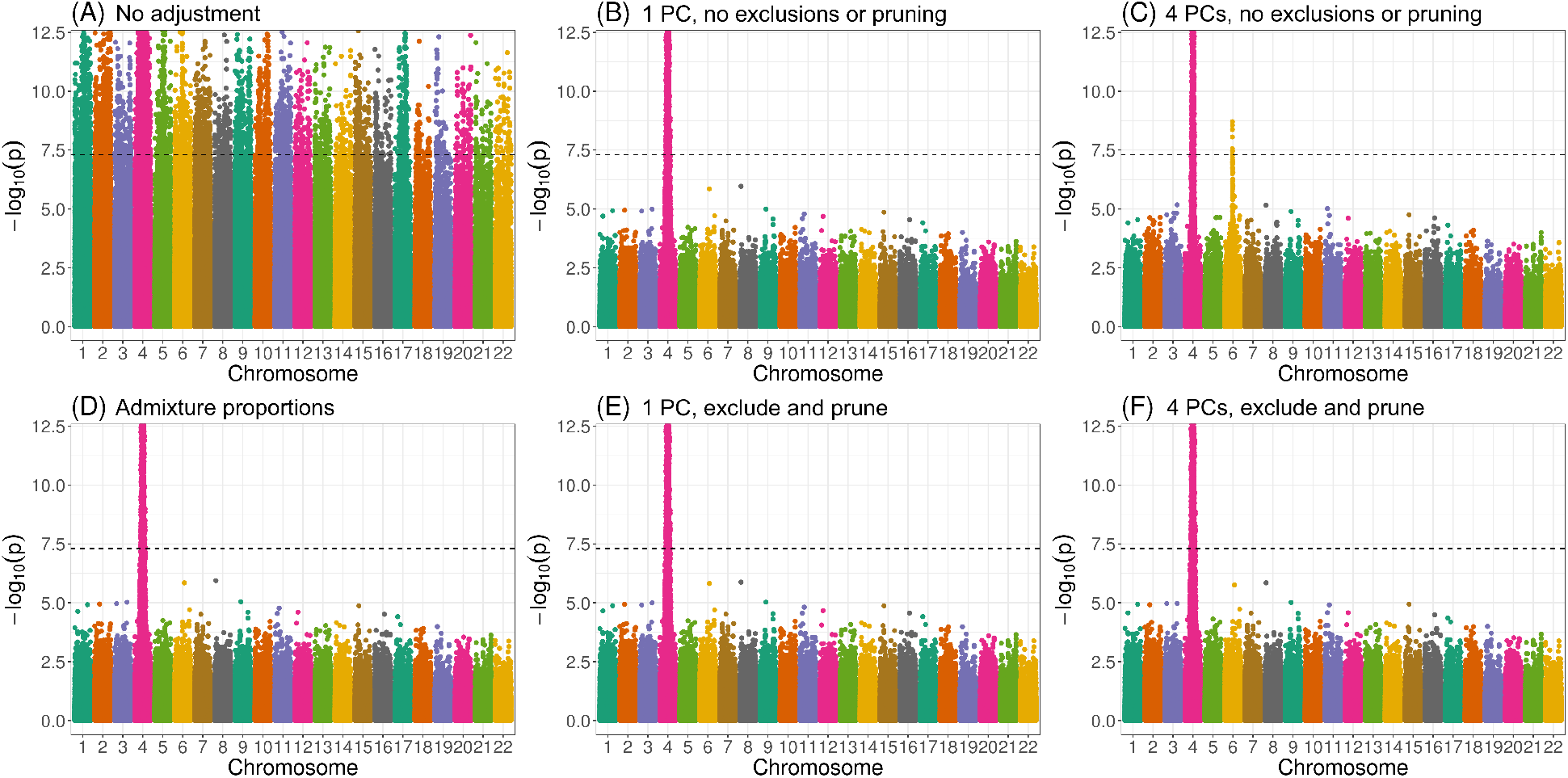
Manhattan plots from GWAS in WHI SHARe African Americans using different approaches to adjust for ancestral heterogeneity. In this example, the simulated trait depends only on the genotype at a single variant on chromosome 4. Panels present results using different adjustment approaches: (A) no adjustment; (B) one PC, with PCs calculated using all variants; (C) four PCs, with PCs calculated using all variants; (D) estimated admixture proportions; (E) one PC, with PCs calculated after LD pruning (*r*^2^ *<* 0.1, window size = 0.5 Mb) and Table 1 exclusions; and (F) four PCs, with PCs calculated after LD pruning and exclusions. The horizontal dashed line in all panels represents the genome-wide significance threshold of 5 × 10*^−^*^8^.

These results are not unique to the choice of causal variant. In Figure 5, we see that adjusting for PCs that captured local genomic features led to higher numbers of spurious associations, on average, across all settings of our simulation using WHI SHARe data. Comparing models that made some sort of adjustment for ancestral heterogeneity, we observed the largest number of spurious associations when GWAS models adjusted for four PCs without any prior LD-based pruning or exclusions (represented by the orange solid line with circles in Figure 5). Excluding the high LD regions from Table 1 prior to running PCA (the orange solid line with triangles) reduced the number of observed spurious associations slightly, but not to the levels of the other approaches. Given what we saw in Figure 3, this is not surprising: even with these exclusions, PCs 2–4 still captured local genomic features— unless those exclusions were also combined with strict LD pruning. On the other hand, when models only included PCs that did not capture local genomic features, the rate of spurious associations dropped. This includes models that adjusted for just the first PC (the green lines in Figure 5) and models that included four PCs, but only after strict LD pruning (the orange dashed lines). Models that adjusted for estimated admixture proportions (the purple solid line) performed nearly identically to models that adjusted for the first PC. This, again, is not surprising, given the high correlation between admixture proportions and the first PC observed in this sample (Supplemental Figures S12 and S13).

**Figure 5:**
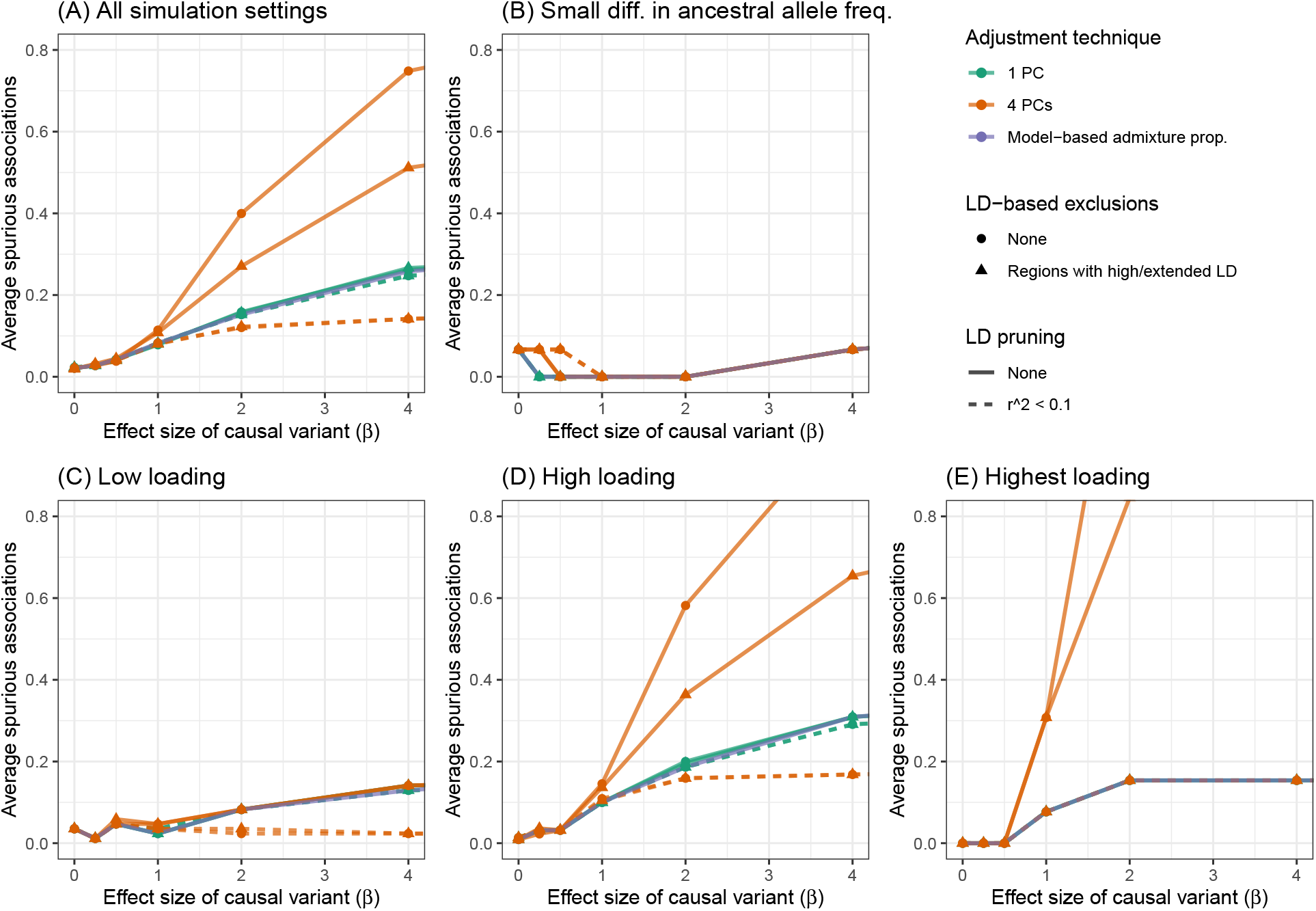
Comparison of the number of spurious associations in GWAS in WHI SHARe African Americans using different approaches to adjust for ancestral heterogeneity. Panel (A) displays the average number of spurious associations that were observed across all simulation settings. Remaining panels focus on the subset of simulation settings in which the causal variant has (B) a small difference in ancestral allele frequencies, (C) low SNP loadings for each of the first four PCs, (D) a high SNP loading for at least one of the first four PCs, or (E) the highest SNP loading on its chromosome for one of the first four PCs. Within each panel, we compare the number of spurious associations when GWAS models adjust for estimated admixture proportions, 1 PC (with or without LD pruning and/or Table 1 exclusions), or 4 PCs (with or without LD pruning and/or Table 1 exclusions). Results shown here are for simulated traits with a single causal variant with effect size (*β*) ranging from 0 to 4.

### 2.6 Factors that influence the rate of spurious associations

Our simulation results highlight various factors that influence when, and how many, spurious associations arise when adjusting for PCs that capture local genomic features. First, we note that there were very few spurious associations, regardless of the adjustment approach, when there were small differences in ancestral allele frequencies at the causal variant (Figure 5B). Even models that made no adjustment for ancestry whatsoever performed similarly in this setting (Supplemental Figure S18). This is to be expected: in this scenario, the causal variant was not associated with global ancestry, so global ancestry was not a confounding variable and thus adjustment was not needed. Considering other simulation settings in which the causal variant had a larger difference in ancestral allele frequencies (panels C, D, and E of Figure 5), and thus adjusting for ancestral heterogeneity was needed, the number of observed spurious associations remained low for models that adjusted for admixture proportions, a single PC (regardless of pre-processing), or four PCs—if those PCs were generated after strict LD pruning. For the two models that adjusted for PCs capturing local genomic features (i.e., the models that adjusted for 4 PCs that were generated with or without Table 1 exclusions, but no LD pruning), however, we saw a higher rate of spurious associations, particularly when the causal variant was highly correlated with one of those PCs. Notably, as the size of the causal variant’s SNP loading increased from low (Figure 5C), to high (Figure 5D), to the highest on its chromosome (Figure 5E), we saw an increasing number of spurious associations for these two approaches. This mirrors the pattern we saw in Figure 4, where a spurious association arose when we adjusted for PCs that were highly correlated with variants in several regions across the genome, and both the causal variant and spurious signal were located in one of those regions. Finally, we note that these problems worsened as the effect size of the causal variant increased.

#### 2.6.1 Connecting simulation results to theory

The patterns observed in our simulation studies are in agreement with our work to derive the expected effect size estimates for GWAS models using different techniques to adjust for ancestral heterogeneity. The setup, assumptions and derivations for this theoretical work are presented in Sections 4.3.2 and S4. In brief, we derived the expected effect size estimate (*E*[*β̂*]) for each model and compared this to the true effect size (*β*). These derivations shed light on the situations in which GWAS models yield biased estimates and the factors that impact the magnitude of this bias.

Figure 6 summarizes our findings about GWAS models that do not make any adjustment for ancestral heterogeneity. We see that these unadjusted models yield a biased estimate of the effect size of the causal variant (*E*[*β̂*] ≠ *β*_1_]) unless there is no ancestral heterogeneity (i.e., Var(*π*) = 0), global ancestry does not have a direct effect on the trait (i.e., *β_π_* = 0), or the causal variant has the same allele frequencies in the two ancestral populations (i.e., *p*_11_ = *p*_10_). We see, also, that the unadjusted model can yield a biased effect size at the unlinked neutral variant (*E*[*β̂*_2_] ≠ 0) even if global ancestry does not have a direct effect on the trait, provided that there is ancestral heterogeneity (Var(*π*) *>* 0) and both the causal variant and the variant being tested have allele frequencies that differ between the two ancestral populations (i.e., *p*_11_ ≠ *p*_10_ and *p*_21_ ≠ *p*_20_). These biased effect size estimates at neutral variants will translate into spurious associations as sample sizes increase, just as we saw in our simulations when we used models that did not adjust for ancestral heterogeneity in any way (Figure 4A and Supplemental Figure S18).

**Figure 6:**
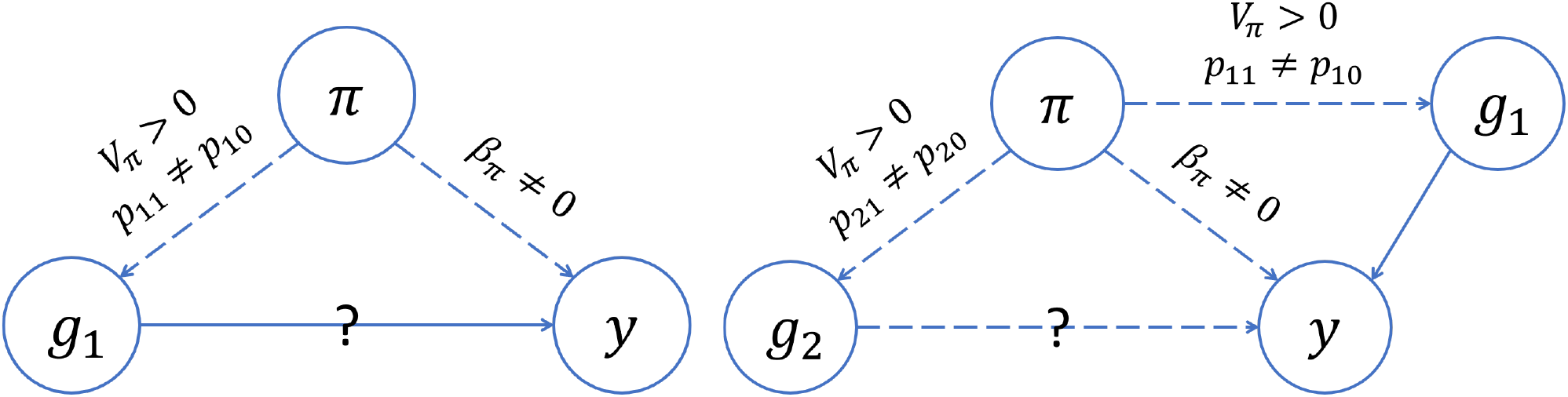
Directed acyclic graphs (DAGs) summarizing the conditions for confounding by global ancestry in GWAS. Each circle represents a variable: *y* is the trait; *π* is global ancestry; *g*_1_ is the genotype of Variant 1, the causal variant; *g*_2_ is the genotype of Variant 2, a second variant that is not associated with the trait and is unlinked with Variant 1. Dashed and solid lines represent relationships between variables, with arrows indicating the direction of the relationship. We used a solid line to indicate a relationship we assume to be present and a dashed line to indicate a relationship that may be present depending on whether certain conditions are met; those conditions are indicated by the text above the line. The question marks indicate the relationships we are interested in testing. On the left, we see that global ancestry confounds the association at the causal variant (Variant 1) if there is ancestral heterogeneity in the population (*V_π_ >* 0), the causal variant has different allele frequencies in the ancestral populations (*p*_11_ ≠ *p*_10_), and global ancestry has a direct effect on the trait (*β_π_* ≠ 0). On the right, we see that global ancestry can confound the association at an unlinked neutral variant (Variant 2) even if global ancestry does not have a direct effect on the trait (*β_π_* = 0), provided that there is ancestral heterogeneity (*V_π_ >* 0) and both the causal variant and the variant being tested have different allele frequencies in the ancestral population (*p*_11_ ≠ *p*_10_, *p*_21_ ≠ *p*_20_).

In contrast, our theoretical work shows that models that appropriately adjust for ancestral heterogeneity will yield unbiased estimates of the effect size at the causal and unlinked neutral variants (Supplemental Equations S3 and S4). At the causal variant, the same holds true even when models adjust for an extraneous PC (Supplemental Equation S5). However, at the unlinked neutral variant these same models will see effect size estimates that are biased away from zero when there is ancestral heterogeneity (i.e., Var(*π*) *>* 0) and the model includes an extraneous PC that is correlated with both the causal variant and the variant being tested (i.e., Cov(*g*_1_*, z* | *π*) ≠ 0 and Cov(*g*_2_*, z* | *π*) ≠ 0)) rather than global ancestry. In other words, these results indicate that if a model adjusts for a PC that is correlated with the causal variant as well as a second variant that is not associated with the trait, then spurious associations will arise at that second neutral variant in large enough samples. This is exactly what we observed in our simulations (Figure 4C, Figure 5D, Figure 5E). However, if the extra PC is not correlated with the causal variant, then spurious associations will not arise; again, this is in agreement with our simulation results (Figure 4F, Figure 5C).

## 3 Discussion

In this paper, we compared approaches for adjusting for ancestral heterogeneity in GWAS in admixed populations, with a particular focus on PCA and its associated pre-processing steps. We showed that PCs can capture local genomic features instead of global ancestry, and that GWAS models adjusting for these PCs have biased effect size estimates and elevated rates of spurious associations. Following recommendations from prior studies, we investigated the impact of removing known high LD regions (Table 1) and LD pruning prior to constructing PCs. We found that the former is not effective in preventing PCs from capturing local genomic features or in reducing the spurious associations that arise as a result. LD pruning proved more effective,although it may not provide a universal solution. Ideal LD parameter settings varied across the datasets investigated in this study, and in one case PCs continued to capture local genomic features even after strict LD pruning. Using model-based estimates of global ancestry, on the other hand, effectively controlled for ancestral heterogeneity in all of our simulations and thus may provide an attractive alternative to PCA in settings where the correlation between PCs and small regions of the genome cannot be easily eliminated. Altogether, our work reiterates the importance of adjusting for ancestral heterogeneity in GWAS in admixed populations and the need for careful consideration of the techniques used to make such an adjustment.

We observed considerable variability in global ancestry proportions across all three admixed populations studied in this paper: the Women’s Health Initiative SNP Health Association Resource (WHI SHARe), Trans-Omics for Precision Medicine Jackson Heart Study (TOPMed JHS), and TOPMed Genetic Epidemiology of Chronic Obstructive Pulmonary Disease Study (COPDGene) African Americans. It is widely understood that adjusting for this ancestral heterogeneity in GWAS is needed in order to control for potential confounding by global ancestry and the spurious associations that can arise as a result. As we showed above, this confounding can occur even when global ancestry does not have a direct effect on the trait itself, provided that there is a causal variant elsewhere in the genome that has different allele frequencies across the ancestral populations of interest. Although this fact has been recognized previously (e.g., [68]), it is sometimes overlooked. Our theoretical work (Supplemental Equations S1 and S2) illustrates the factors that impact the magnitude of the bias incurred by GWAS models that fail to adjust for global ancestry. We hope that our results will serve as a reminder to researchers of the various ways in which global ancestry can confound genetic studies in admixed populations and the importance of ensuring that GWAS models appropriately adjust for ancestral heterogeneity.

A common approach for adjusting for ancestral heterogeneity in GWAS involves including global ancestry as a covariate in marginal regression models, with global ancestry estimated using either model-based approaches or PCA. In WHI SHARe, TOPMed JHS, and TOPMed COPDGene African Americans, the first PC was highly correlated with estimates of the genome-wide proportion of African ancestry and models adjusting for either performed similarly. Later PCs, however, did not correlate with global ancestry and models that adjusted for these PCs anyway—a common practice in the literature—yielded elevated rates of spurious associations, particularly when those PCs captured local genomic features rather than genome-wide ancestry. Prior work (e.g., [54,56]) has shown that PCs can detect regions with high, extensive, or otherwise unusual patterns of LD (Table 1), but the patterns observed in those studies—primarily involving individuals of European ancestry—differ from what we saw here. In particular, those studies typically saw PCs that were driven by variants in a single high-LD region, just as we saw in TOPMed COPDGene European Americans (Supplemental Figure S9). However, in all three of the admixed samples that we investigated, we saw instead that PCs were correlated with variants in *multiple* regions, across multiple chromosomes. This has important downstream implications.

Our work shows that adjusting for PCs that capture multiple local genomic features can induce spurious associations. This issue can be explained by the concept of *collider bias* (Figure 7). If a PC is correlated with the genotype of multiple variants, and at least one of those variants is associated with the trait, then the PC becomes a *collider variable* when testing the association between the other variants and the trait. Adjusting for collider variables can induce a spurious association between variables that are otherwise unlinked [69]. This is precisely what we saw above. GWAS models adjusting for an extraneous PC will yield biased estimates of variant effect sizes, with the magnitude of that bias increasing with the effect size of the causal variant, the strength of the correlation between the PC and the variants it captures, and the amount of ancestral heterogeneity within the population (Supplemental Equation S6). When that bias is large enough, it can lead to a spurious association—as our simulations show. Given the similarities in terms of which regions of the genome tend to be correlated with PCs across different admixed populations (see Figures 2 and 3, for example), it is possible that these spurious associations may even replicate across studies.

**Figure 7:**
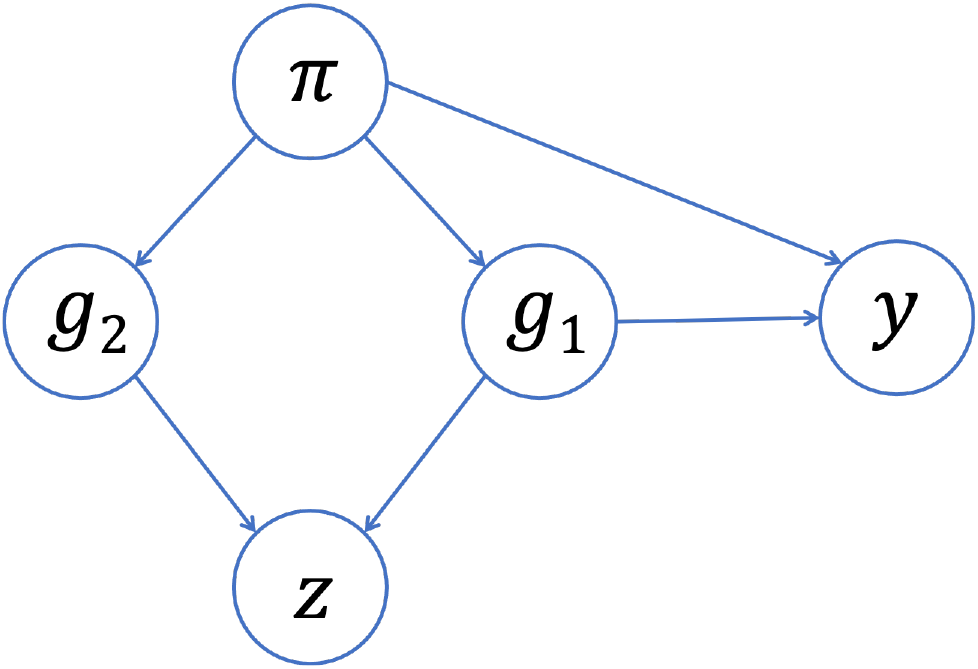
Collider bias in GWAS. Suppose that, instead of genome-wide global ancestry (*π*), a PC (*z*) captures the genotype of two variants (*g*_1_, *g*_2_). If one of those variants (*g*_1_) is associated with the trait (*y*), then the PC will be a collider variable—rather than a confounding variable—when testing the association between the other variant (*g*_2_) and the trait. Adjusting for the PC can then induce a spurious association between the trait and the neutral variant as a result of collider bias.

The focus in the literature has typically been on demonstrating issues that can arise when adjusting for *too few* PCs [7, 18, 48, 49, 70]. Prior discussion of the impact of adjusting for PCs capturing local genomic features have focused on persistent inflation due to sub-optimal ascertainment of population structure [45, 47, 50, 54, 55, 60, 61] and decreased power when a causal variant is located in one of the regions highly correlated with a PC [45, 47, 49, 54–56, 62, 63]. In studying the impact on type I error, many of these studies focus on the inflation factor *λ* [45, 50, 60, 62]. Although useful for detecting widespread inflation, our simulations suggest that inflation factors are not sensitive to the presence of spurious associations arising from collider bias (Supplemental Information Section S5.2). This could partially explain why the issue has not been detected previously. Furthermore, the vast majority of these studies focused on European populations, where PCs are typically correlated with variants on just a single chromosome (e.g., Supplemental Figure S9) and thus collider bias is not a concern. Those studies that did see multiple peaks in SNP loading plots (e.g., [54, 56, 60]) did not make any mention of its implications. We did identify one prior study [49] that investigated the performance of PCA in a simulated admixed population; they saw slightly higher type I error rates when using 10 PCs instead of 2 (consistent with our findings), but the authors did not provide any discussion of what those later PCs might have captured or what may have caused the elevated type I error rate.

In recent years, increasing attention has been paid to the issue of collider bias in genetic association studies [71–74], but to the best of our knowledge this is the first paper to fully demonstrate the concerns related to including extraneous PCs in GWAS models. Most closely related to our work, Dahl et al. [75] showed that adjusting for PCs can lead to replicating spurious associations in gene expression studies due to collider bias. As a short aside, the authors mentioned that collider bias could arise in GWAS, but they implied that the magnitude of this bias would be small— and thus not of practical concern—given that “causal SNPs have much lower leverage on genetic PCs” (i.e., the correlation between PCs and causal variants is small). Our findings are consistent with their conclusion that the magnitude of collider bias depends on the strength of the correlation between the PCs and the variants they capture. However, we found that this correlation between PCs and genetic variants can be non-trivial in admixed populations (without careful pre-processing prior to running PCA), implying that collider bias is of more concern in this setting.We also showed that the amount of bias depends on the variance of admixture proportions across a population (denoted *V_π_* in Supplemental Equation S6), which could explain why work that has focused on more ancestrally homogenous populations may not have identified this issue previously. That said, it is important to raise the question of the magnitude of bias that could be expected in a “typical” GWAS. The smaller the effect size of the causal variant, the smaller the number of spurious associations that we observed in our simulation study (Figure 5), and when studying complex traits/diseases we would expect the causal variant effect sizes to be fairly small. However, in these settings we would also expect that there are *multiple* causal variants—not just a single causal variant, as we assumed for the sake of simplicity in our simulations and theoretical work—and then these effects may be aggregated [76]. In any case, it is worth considering what steps can be taken to reduce or eliminate concerns about potential collider bias altogether.

In our analysis of genotype and sequence data from unrelated WHI SHARe, TOPMed JHS, and TOPMed COPDGene African Americans, we found that all but the first PC were correlated with small regions of the genome—and thus have the potential to be collider variables—unless careful pre-processing of genotype data was performed prior to running PCA. As mentioned earlier, previous studies have found that PCs can be driven primarily by small regions of the genome, and as a result have suggested that these regions be excluded (Table 1) and/or that LD pruning be performed prior to running PCA. However, the motivation for this LD-based filtering has typically been framed in terms of the ability of the PCs to capture global ancestry, as well as the computational complexity of running PCA, rather than the downstream implications on association testing that we have highlighted here. Furthermore, we found that excluding the regions listed in Table 1, without also performing LD pruning, did not solve these issues in admixed populations: we still saw peaks in the PC-genotype correlation plots and models adjusting for these PCs continued to show elevated rates of spurious associations. A more tedious iterative approach of identifying and removing potentially problematic regions based on our own data (similar to that proposed by [61]) also did not prevent PCs from capturing local genomic features within a reasonable number of iterations (Supplemental Figure S3). LD pruning, on the other hand, proved to be more successful, at least when looking at the first four PCs in WHI SHARe African Americans. Our results highlight that a stricter threshold (e.g., *r*^2^ = 0.1) may be needed for LD pruning in admixed populations than the *r*^2^ = 0.2 threshold that is often suggested in the literature: see Supplemental Information Section S1.1 for a comparison. Given that LD patterns differ between admixed populations and the European populations upon which much of this prior work was based, it is not surprising that different LD-based filtering techniques are required here.

It is important to acknowledge, however, that LD pruning with the parameters we used in WHI SHARe African Americans (*r*^2^ = 0.1, window size = 0.5 Mb) may not be a universal solution. For example, in TOPMed JHS African Americans, LD pruning with these parameters partially, but not completely, resolved the issue of PCs being highly correlated with variants on multiple chromosomes. Comparing Figure 2A to Supplemental Figure S4, we saw improvement for the second and third PCs, but there continued to be some peaks (albeit shorter) in the PC-genotype correlation plot for the fourth PC. It was not until we used a wider window size of 10 Mb that the peaks disappeared (Supplemental Figure S5) for all of the top four PCs. In TOPMed COPDGene African Americans, even this wider window size was not sufficient in preventing PCs from capturing local genomic features (Supplemental Figures S7).

Our analyses in this paper have focused on three samples of African American individuals, but we believe that our findings generalize more broadly. We have, for example, observed similar patterns of PCs capturing multiple local genomic features through our involvement in a study including African American, African Caribbean, and Hispanic/Latino TOPMed participants [77], as well as a study focused on individuals of Mexican descent [78]. That said, we have not conducted an exhaustive study of all admixed populations. It is also important to acknowledge that patterns, implications, and ideal solutions may differ in other non-European populations or in trans-ancestry analyses. Thus in any sample—admixed or otherwise—we strongly recommend that investigators check SNP loadings or the correlation between PCs and genotypes before including PCs in GWAS models. If multiple variants are associated with a PC and sample size is sufficiently large, spurious associations can arise. In some samples, it may be necessary to consider stricter LD pruning (e.g., smaller *r*^2^ threshold, larger window size), to include fewer PCs, or to consider an alternate approach altogether (e.g., adjusting for estimated admixture proportions).

Ultimately, our work demonstrates the challenges that can arise in appropriately adjusting for ancestral heterogeneity in admixed populations. Compared to prior work in European populations, we see that the patterns of which and how many local genomic features are captured by PCs differs, leading to distinct downstream implications for GWAS, namely, collider bias. The previously recommended pre-processing step of removing known high LD regions (Table 1) shows limited utility in admixed populations, and the effectiveness of LD pruning is sensitive to the choice of parameters. These results are particularly concerning given the wide-spread use of PCA in the GWAS literature, and the inconsistent use and reporting of pre-processing steps. For populations where we have a good idea of the number of ancestral populations of interest and relevant reference panel data is readily available, GWAS models adjusting for estimated global ancestry proportions rather than PCs perform well. PCA can offer advantages over model-based ancestry inference methods, but careful consideration must be given to how many PCs should be included and what those PCs are capturing. In the African American samples studied here, for example, a single PC was sufficient for controlling spurious associations induced by population structure. In some simulation settings, we did see a small drop in the number of spurious associations when we included three additional PCs, but only after strict LD pruning. Careful pre-processing of data prior to running PCA, combined with thorough diagnostics (i.e., calculating and plotting SNP loadings or the correlation between PCs and genotypes, rather than simply relying on inflation factors), is critical. We urge investigators to carefully check PCs before including them in GWAS models, to clearly justify their choice of which and how many PCs to include, and to transparently report the steps they have taken to ensure that those PCs are not causing the very problem—spurious associations—that these techniques aim to solve.

## 4 Materials and Methods

### 4.1 Data and Quality Control

Our analyses focus on array-based and whole genome sequence-based genotype data from three samples of African American individuals. In particular, we consider genotype data from the Women’s Health Initiative SNP Health Association Resource (WHI SHARe), as well as whole genome sequencing data from two contributing studies to the Trans-Omics for Precision Medicine (TOPMed) Whole Genome Sequencing Project: the Jackson Heart Study (JHS) and the Genetic Epidemiology of Chronic Obstructive Pulmonary Disease Study (COPDGene). We performed quality control and identified subsets of unrelated African American individuals prior to running any further analyses—details are provided below.

#### 4.1.1 WHI SHARe Genotype Data

The Women’s Health Initiative (WHI) is a long-term study of the health of post-menopausal women residing in the United States. In total, 161,808 women aged 50–79 years old were recruited to participate in this study. Additional details of the study design and cohort characteristics can be found elsewhere [79]. The WHI SHARe study includes 12,151 self-identified African American women who consented to genetic research, a subsample of which were selected for genotyping using the Affymetrix Genome-Wide Human SNP Array 6.0. This array contains 906,000 single nucleotide polymorphisms (SNPs) and more than 946,000 probes for detection of copy number variants. In our analyses, we focus only on the SNP data. We did not impute WHI genotypes beyond filling in sporadic missing genotypes—see Grinde et al. [80] for more details.

The genotype data were processed for quality control. After filtering on call rate, concordance rates for blinded and unblinded duplicates, and sex discrepancy, there were 871,309 SNPs with a missing genotyping rate of 0.2% and 8,421 African American women [24]. We also used the iterative procedure suggested by Conomos et al. [81] to identify a subset of 8,064 mutually unrelated individuals, using a kinship threshold of 0.044 (i.e., excluding first-, second-, and third-degree relatives).

#### 4.1.2 TOPMed Whole Genome Sequence Data

The TOPMed Whole Genome Sequencing Project is an ongoing project sponsored by the National Heart, Lung, and Blood Institute. The goal of this project is to collect and analyze whole-genome sequences, other omics data, and extensive phenotypic information for over 100,000 individuals from diverse backgrounds. Data are periodically released on dbGaP for analysis by the broader scientific community. Our analysis uses data from freeze 4, released in 2017, and freeze 5b, released in 2018. These two freezes include samples from a large number of contributing studies. We focus on two of these studies: the Jackson Heart Study (JHS) (accession number: phs000964) and the Genetic Epidemiology of Chronic Obstructive Pulmonary Disease Study (COPDGene) (accession number: phs000951). High coverage (≈ 30X) whole genome sequencing was performed by the University of Washington (UW) Northwest Genomics Center for JHS (freeze 4) and by a combination of the UW Northwest Genomics Center and the Broad Institute of MIT and Harvard for freeze 5b COPDGene. Details on TOPMed sequencing and QC methods are available in Taliun et al. [82] and on the TOPMed website: https://topmed.nhlbi.nih.gov/data-sets. In total, the downloaded freeze 4 JHS dataset includes 2,777 African American individuals and the freeze 5b COPDGene dataset includes 8,476 African American and European American individuals consented for biomedical research.

Prior to genetic ancestry inference, we performed two additional stages of variant- and sample-level filtering. We used bcftools [83] to restrict our analyses to biallelic single nucleotide variants (SNVs). To identify a subset of mutually unrelated individuals (kinship threshold = 0.044), we used the UW Genetic Analysis Center (GAC) TOPMed analysis pipeline. In addition to inferring relatedness using the procedure proposed by Conomos et al. [81], this pipeline also includes code to perform PCA, association testing, and other tasks in whole genome sequence data: more details can be found at https://github.com/UW-GAC/analysispipeline. After variant filtering and removal of related individuals, 1,928 and 8,406 unrelated samples and 77,136,850 and 135,522,041 variants remained in JHS and COPDGene, respectively.

### 4.2 Genetic Ancestry Inference

We consider two approaches to inferring genetic ancestry in these admixed samples: model-based approaches and PCA.

#### 4.2.1 Model-Based Approaches

In WHI SHARe African Americans, we inferred both local and global genetic ancestry using model-based ancestry inference techniques. Local ancestry inference was performed using RFMix [32] and a reference panel composed of samples from the CEU (Utah residents with Northern and Western European ancestry) and YRI (Yoruba in Ibadan, Nigeria) populations from the International HapMap Project (HapMap) [67]: see Grinde et al. [80] for more details. We then calculated global ancestry proportions via the genome-wide average local ancestry 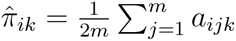, where *a_ijk_* is the inferred number of alleles (0, 1, or 2) inherited by individual *i* at variant *j* from ancestral population *k*. We also compared these RFMix-based global ancestry estimates to results from supervised and unsupervised ADMIXTURE [31] analyses with two ancestral populations (*K* = 2). The supervised analysis used the same HapMap reference panel as was used to infer local ancestry using RFMix. All three sets of admixture proportions were highly correlated (pairwise Pearson correlation *>* 0.998). We proceed with using only the RFMix-based admixture proportion estimates for the remainder of our analyses.

In TOPMed JHS and COPDGene samples, we inferred global ancestry via unsupervised ADMIXTURE analyses with two ancestral populations. We also used these inferred global ancestry proportions to identify subsets of admixed individuals. Although JHS recruited only self-identified African Americans, we did identify 40 individuals in the sample with an estimated European ancestry proportion of 100%. These individuals were excluded from further analyses, leaving a total of 1,888 unrelated admixed samples. The COPDGene study, by design, includes both African American and European American individuals. Self-identified race/ethnicity information was not available from dbGaP, so we used inferred admixture proportions to identify and restrict our attention to individuals with at least 29.5% African ancestry. The choice of threshold follows from the results reported by Parker et al. [84], showing that the self-identified African American individuals in the COPDGene study have inferred proportions of African ancestry ranging from 29.5% and above. (We are not suggesting that this same threshold be universally applied to identify African American individuals in other samples.) After filtering, 2,676 admixed individuals remain in COPDGene. The remainder of our TOPMed analyses focused on these subsets of unrelated admixed individuals.

#### 4.2.2 Principal Component Analysis

To infer global ancestry using PCA, we perform a singular value decomposition of the matrix of standardized genotypes (i.e., **X** = **UDV***^T^*) or, equivalently, an eigenvalue decomposition of the genetic relationship matrix (i.e., **XX***^T^* = **UD**^2^**U***^T^*), where **X** is the *n* × *m* matrix of standardized genotypes for *n* individuals at *m* single nucleotide variants. One or more of the top eigenvectors, or PCs, **u**_1_, **u**_2_*, . . .*, typically reflect global ancestry.

We ran PCA on the WHI SHARe genotype data using SNPRelate [42]. First, we applied PCA to the same set of 551,025 SNPs used to estimate global ancestry proportions. We then applied PCA to subsets of SNPs based on the following pre-processing criteria: excluding SNPs falling into regions of the genome that have been cited in the literature as potentially problematic for PCA (Table 1), LD pruning, or both literature-based exclusions and LD pruning. To perform LD pruning, two parameters must be specified: *r*^2^ threshold and window size. Here we use an *r*^2^ threshold of 0.1 and window size of 0.5 mega basepairs (Mb), which is stricter than is often suggested in the literature: for a full discussion of these choices, see Supplemental Information Section S1.1. Both LD pruning and filtering of regions in Table 1 were implemented using the SNPRelate package. Table 2 summarizes the number of SNPs remaining after each set of pre-processing steps. After running PCA on these different sets of variants, we also used SNPRelate to assess the contribution of each SNP to each PC by calculating the SNP loadings and the correlation between PCs and genotypes.

In TOPMed JHS and COPDGene samples, we used the UW GAC TOPMed analysis pipeline to implement pre-processing, run PCA, and calculate and visualize the contribution of individual variants to each PC. Similar to WHI SHARe, we applied PCA to various subsets of variants based on different pre-processing criteria, including a naive analysis with no prior LD-based pruning or filtering, an analysis after excluding the regions listed in Table 1, an analysis after LD pruning with an *r*^2^ threshold of 0.1, and an analysis after both LD-based pruning and filtering. Following the recommendations of Kirk [85] and the UW GAC pipeline documentation, we also removed variants with a minor allele frequency lower than 0.01 in all cases. Note that the UW GAC pipeline provides the option to exclude some of the regions listed in Table 1 (the *LCT* gene on chromosome 2, the HLA region on chromosome 6, and the locations of large inversions on chromosomes 8 and 17), but we customized the pipeline code to add the other regions identified in our literature review. See Table 2 for the number of variants included in each analysis.

### 4.3 Investigation of Downstream Implications for GWAS

To explore the impact of adjusting for PCs that capture local genomic features, as well as to further evaluate the utility of the proposed solutions of LD pruning and removing known high LD regions (Table 1) prior to running PCA, we derived expected effect size estimates and conducted simulation studies comparing the rate of spurious associations for GWAS models adjusting for different sets of covariates.

#### 4.3.1 GWAS Models

We consider models of the general form

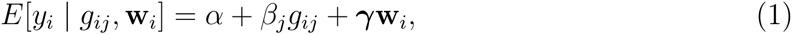

where *y_i_*is the quantitative trait, *g_ij_*is the genotype at position *j*, and **w***_i_* is a vector of additional covariates. Note that we quantify genotype *g_ij_* by the number of copies—0, 1, or, 2—of some pre-specified allele (e.g., the minor allele) carried by individual *i* at position *j*. We fit these models at every position *j* = 1*, . . ., m* across the genome and test for association between the trait and genotype by testing the null hypothesis *H*_0_ : *β_j_* = 0 using a traditional Wald test.

In particular, we are interested in comparing four models: a model making no adjustment for ancestral heterogeneity (i.e., **w***_i_* = ∅ø), a model adjusting for estimated admixture proportions (**w***_i_* = *π̂_i_*), a model adjusting for the first PC (**w***_i_* = *u*_1_*_i_*), and models adjusting for additional PCs (e.g., 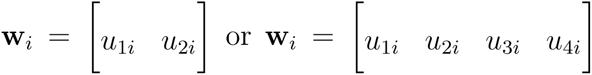). For the models adjusting for PCs, we consider four sets of PCs based on different pre-processing criteria: *none* (no prior LD-based exclusions or pruning), *exclusions only* (excluding regions from Table 1 but no LD pruning), *pruning only* (LD pruning with *r*^2^ *<* 0.1 and a window size of 0.5 Mb, but not excluding regions from Table 1), and *both* (both Table 1 exclusions and LD pruning).

#### 4.3.2 Effect Size Derivations

In deriving expected GWAS effect size estimates, we assume that the admixed population has two ancestral populations and the trait depends on a single causal variant:

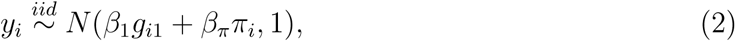

where *g_i_*_1_ represents the number of minor alleles carried by individual *i* at the causal variant, which we will refer to as *Variant 1*, and *π_i_*is the individual’s admixture proportion. The coefficients *β*_1_ and *β_π_*represent the effect of the causal variant and global ancestry, respectively, on the trait. More details of the assumed data generating mechanism can be found in Supplementary Information Section S4.

Following the framework outlined above, we compare models that adjust for the true admixture proportions (i.e., Model 1 with **w***_i_* = *π_i_*) and models that does not make any adjustment for ancestral heterogeneity (i.e., Model 1 with **w***_i_* = ∅ø). To mimic the scenario of adjusting for extraneous PCs, we also consider models that adjusts for two “PCs” where the first “PC” captures global ancestry but the second “PC” captures some feature other than global ancestry (i.e., Model 1 with 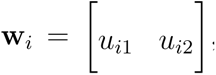, where **u**_1_ = ***π*** and **u**_2_ = **z** for some variable *z*). For each of these models, we derive the expected effect size estimate at the causal variant (Variant 1), as well as a second variant that is not associated with the trait and sits on a different chromosome than the causal variant. Detailed results are presented in Supplementary Information Section S4.

#### 4.3.3 Simulation Studies

We conducted simulation studies to support and expand upon the findings from our effect size derivations described above. Here, we focus in particular on simulations that highlight the impact of adjusting for PCs that capture local genomic features and investigate whether including extraneous PCs can induce spurious associations.

In WHI SHARe, we simulated traits for each individual *i* = 1*, . . .,* 8064 such that they depended only on the genotype *g_ij_* at a single causal variant with effect size *β_j_* (Equation 2 with *β_π_* = 0). We considered various choices of the effect size (*β_j_* = 0, 0.25, 0.5, 1, 2, 4, 8) and nearly 500 choices for the position *j* of the causal variant, with the goal of investigating the impact of the effect size of the causal variant *β_j_*, the difference in its ancestral allele frequencies, and the strength of its contribution to top PCs—all of which our theoretical results (Supplementary Information Section S4) suggest contribute to the magnitude of the bias observed in effect size estimates from GWAS models adjusting for PCs that capture local genomic features.

To choose these causal variants, we first estimated the difference in ancestral allele frequencies for each variant using the observed allele frequencies in our HapMap reference panel (see Section 4.2.1). We also considered the SNP loadings for the set of PCs that were generated without any prior LD-based filtering or pruning (Section 4.2.2). For each of the first four PCs, we identified the 10 variants on each chromosome with the highest absolute SNP loadings. For comparison, we also selected 100 variants across the autosomes with low SNP loadings (|loading| *<* 0.0008) for all of the first four PCs. We chose these variants such that most (85) had substantially different allele frequencies in the African and European ancestral populations (|*p̂_CEU_* − *p̂_Y RI_*| *>* 0.6), while others (15) had similar allele frequencies in the two ancestral populations (|*p̂_CEU_* − *p̂_Y RI_*| *<* 0.005). Altogether, this amounted to 473 causal variants with positions spread across the genome, high or low SNP loadings, and large or small ancestral allele frequency differences, allowing us to investigate the impact of different characteristics of the causal variant on GWAS results.

For each simulation replicate, we fit GWAS models using a variety of techniques to adjust for ancestral heterogeneity (Section 4.3.1). We evaluated these techniques by comparing the number of spurious associations that appeared when we used each model. We quantified spurious associations by counting the number of chromosomes, not including the chromosome on which the causal variant was located, with at least one variant reaching genome-wide significance. Note that we chose to count spurious associations at the chromosome level rather than the individual variant level because variants near each other on the same chromosome may be in strong LD and will thus show similar association results. The genome-wide significance threshold was set to the *p* = 5.0 × 10*^−^*^8^ threshold that is used extensively throughout the GWAS literature [86, 87].

In TOPMed African American samples, we similarly simulated quantitative traits depending only on the genotype at a single variant. We constructed two “PCs” such that the first was equal to the estimated African admixture proportion (Section 4.2.1) and the second depended on the genotype at the causal variant as well as another variant on a distinct chromosome. We compare the results of GWAS models adjusting for a single PC versus models that also adjust for the extraneous second PC that captures local genomic features. In these simulations, we used a significance threshold of 5.0 × 10*^−^*^9^ following recommendations for association studies in admixed populations using whole genome sequence data [88].

## Acknowledgments

The WHI program is funded by the National Heart, Lung, and Blood Institute, National Institutes of Health, U.S. Department of Health and Human Services through 75N92021D00001, 75N92021D00002, 75N92021D00003, 75N92021D00004, 75N92021D00005. The authors than the WHI investigators and staff for their dedication, and the study participants for making the program possible. A short list of WHI investigators can be found in the Supplemental Information (Section S6), and a full listing of all investigators who have contributed to WHI can be found at https://www.whi.org/doc/WHIInvestigator-Long-List.pdf. Molecular data for the Trans-Omics in Precision Medicine (TOPMed) program was supported by the National Heart, Lung, and Blood Institute (NHLBI). Core support including centralized genomic-read mapping and genotype calling along with variant quality metrics and filtering were provided by the TOPMed Informatics Research Center (3R01HL-117626-02S1; contract HHSN268201800002I). Core support including phenotype harmonization, data management, sample identity QC, and general program coordination were provided by the TOPMed Data Coordinating Center (R01HL-120393; U01HL-120393; contract HHSN268201800001I). We gratefully acknowledge the studies and participants who provided biological samples and data for TOPMed. The Jackson Heart Study is supported and conducted in collaboration with Jackson State University (HHSN268201300049C and HHSN268201300050C), Tougaloo College (HHSN268201300048C), and the University of Mississippi Medical Center (HHSN268201300046C and HHSN268201300047C) contracts from NHLBI and the National Institute for Minority Health and Health Disparities (NIMHD); genome sequencing was funded by HHSN268201100037C. The COPDGene study was supported by NIH grants U01 HL089856 and U01 HL089897. The COPDGene project is also supported by the COPD Foundation through contributions made by an Industry Advisory Board comprised of Pfizer, AstraZeneca, Boehringer Ingelheim, Novartis, and Sunovion.

## Supporting Information

**S1 Text. Supplemental Information.** Includes 15 figures, proofs and simulations validating the theoretical results presented in the main paper, and a list of WHI investigators.

## Data and Code Availability

WHI SHARe genotype data is available on dbGaP (accession number: phs000386) or directly through WHI according to the policy outlined at https://www.whi.org/doc/WHI-genetic-data-transfer-policy.pdf. TOPMed whole genome sequence data is also available from dbGaP (Jackson Heart Study: phs000964, Genetic Epidemiology of Chronic Pulmonary Disease Study: phs000951). All software packages used throughout this paper are freely available online:

- bcftools [83] (quality control): https://samtools.github.io/bcftools/
- RFMix [32] (local ancestry inference): https://sites.google.com/site/rfmixlocalancestry
- ADMIXTURE [31] (global ancestry inference): https://dalexander.github.io/admixtur
- PCRelate [81] and PC-AiR [43] (identifying unrelated individuals): https://rdrr.io/bio
- SNPRelate [42] (LD pruning, PCA, and PCA-related diagnostics): https://www.bioconductor.org/packages/release/bioc/html/SNPRelate.html
- PLINK [64] (GWAS): https://zzz.bwh.harvard.edu/plink/
- TOPMed Analysis Pipeline (identifying unrelated individuals, LD pruning, PCA, PCArelated diagnostics, and association studies in whole genome sequence data): https://github.com/UW-GAC/analysis pipeline
- R (analyzing and visualizing results): https://cran.r-project.org/ Other resources pertaining to this paper, including download-able lists of the high LD regions in Table 1 in various builds, can be found on the lead author’s GitHub page:
- https://github.com/kegrinde/PCA.

